# Fast Assembling of Neuron Fragments in Serial 3D Sections

**DOI:** 10.1101/097253

**Authors:** Hanbo Chen, Daniel M. Iascone, Nuno Macarico da Costa, Ed S. Lein, Tianming Liu, Hanchuan Peng

## Abstract

Reconstructing neurons from 3D image-stacks of serial sections of thick brain tissue is very time-consuming and often becomes a bottleneck in high-throughput brain mapping projects. We developed NeuronStitcher, a software suite for stitching non-overlapping neuron fragments reconstructed in serial 3D image sections. With its efficient algorithm and user-friendly interface, NeuronStitcher has been used successfully to reconstruct very large and complex human and mouse neurons.

---

Digital reconstructions of neurons from very large three-dimensional (3D) brain images is crucial for modern neuroscience^1–3^. Despite recent great advances in neuron labeling, brain clearing, and high-resolution 3D tissue-imaging^4,5^ to study mammalian brains, many neuroscientists still rely on physical sectioning of brains followed by imaging such sections using either light microscopy in 3D or electron microscopy in 2D. The acquired images of serial sections are then stacked and aligned to generate a very big image volume, from which neurons are reconstructed and quantified (e.g. Figure 1(a)-(d)). Dendrites and axons severed at the section boundaries will need to be stitched. This is often a challenging bottleneck for proper reconstruction of locally dense dendritic and axonal trees.

**Figure 1.**
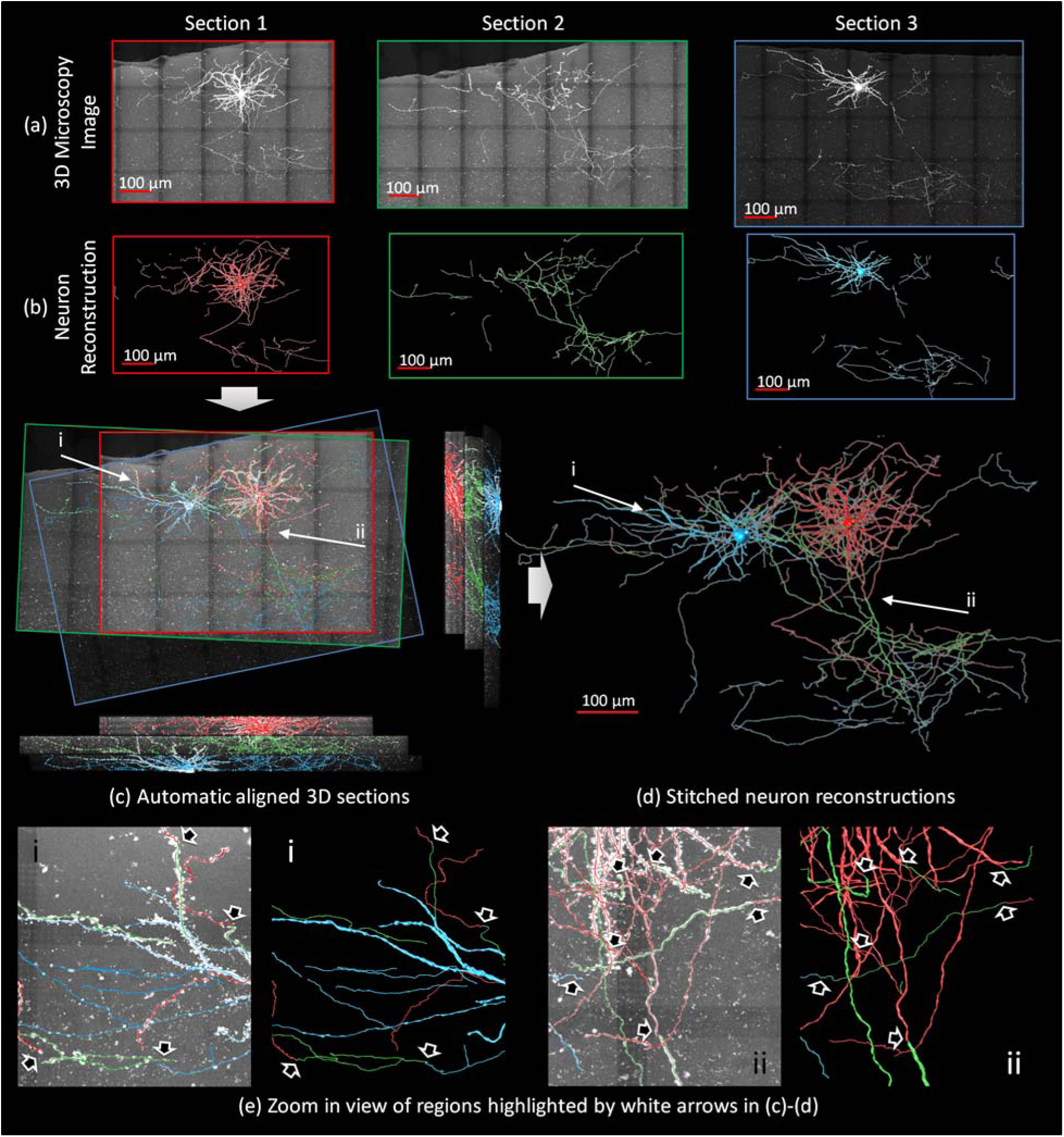
Illustration of the 3-D neuron stitching problem and results. Three continuous sections containing 2 neurons in a mouse brain are shown. The objective was to align neuron reconstructions (b) from adjacent 3D sections (a) and stitch them into complete neuron reconstructions (c)-(d). Different colors (red, green, and blue) were assigned to the reconstructions in each section. All the snapshots were generated from a view perpendicular to the section plane except the two side-view pictures shown on the right and bottom of (c). (e) Zoom-in views of two regions in (c) and (d) with the stitched neurites highlighted by arrows.

It is quite labor-intensive to stitch neuron segments manually over multiple sections. Automated methods may provide a significant increase in the throughput to neuronal reconstruction. However, this is a non-trivial task for algorithms because there could be missing tissues as well as distortions during sectioning, making stitching neuronal segments across multiple sections much more challenging in comparison with stitching overlapping tiles within single sections^6,7^ (Figure 1(c)). Several studies ^8−12^ aimed at aligning traced neuron fragments (Supplement, Section 1) because a neuron reconstruction often gives a concise and less noisy description of neuron morphology than the respective raw image. These methods demonstrated various levels of suitability in identifying an alignment. Yet, most of these tools are not readily available in their Open Source form, thus a thorough evaluation is difficult. Moreover, none of the existing methods was explicitly designed to handle the artifacts such as tissue loss or noise. Hence their performance is sensitive to the quality of neuron reconstruction (Supplement, Section 4).

To address these challenges, we developed NeuronStitcher, a software package that automatically assembles complicated neuron fragments reconstructed from adjacent serial sections. NeuronStitcher utilizes a triangle-matching algorithm to estimate the initial match of severed neurites on the sectioning plane based on their relative location and branch-orientations (Supplement, Section 2.3). Then the reconstruction alignment process is iterated until the final matched neurites are smoothly stitched (Supplement, Section 2.4). To make NeuronStitcher perform well for neurites that have varying quality, we have also developed three methods to remove noise when determining severed neurites (Supplement, Section 2.2).

We evaluated the accuracy of NeuronStitcher using carefully generated “ground truth” reconstructions from a piece of mouse brain tissue containing a labeled pyramidal neuron within the hippocampal CA1 region (Supplement, Section 4.1). Specifically, the whole neuron was first imaged and thereafter semi-automatically reconstructed in 3D by an expert as the ground truth reconstruction (Figure S 10(a)). Then, we evenly sliced the neuron into 3 serial sections in the z direction, each of which was imaged individually. The reconstructions of neuron fragments from all individual sections were generated by an expert and then stitched together using NeuronStitcher (Figure S 10(b)). A careful comparison of the ground truth reconstruction to the stitched reconstruction showed that 98% bifurcations of the 3D reconstructed, tree-like neuron morphology in the ground truth had their correspondence in the stitched reconstruction (Supplement, Section 4.1), and the minor amount (2%) of missing correspondence happened at the section interfaces and was due to the sectioning process.

We also considered an alternative way to produce the “ground truth” to evaluate the accuracy of NeuronStitcher (Supplement, Section 4.3). We chose 5 densely arborized reconstructions from mouse visual cortex. To generate the simulated data, each reconstruction was digitally “sectioned” into two halves. One-half was then randomly rotated and shifted in parallel to the sectioning plane. Several different gaps (1, 2, 4, and 8 μm) were added to simulate the different levels of tissue loss during sectioning. Both the vertices and edges within the sectioning gaps were removed (Figure S 17). To test NeuronStitcher, it was used to stitch the two portions of data back together. Our automatic matching found the correct matching and alignment in most cases (Figure S 18). Even when a considerable amount of tissue was lost (8 μm gap), the alignment was still close to the ground truth (10.4 μm). In such a case, most of the severed neurites were correctly matched (75% precision, 62% sensitivity) by using the default parameters. Notably, the errors due to the substantial information loss in the big gaps should be anticipated (Figure S 18(c)-(e)). We also tested NeuronStitcher with different parameter configurations, which showed that NeuronStitcher was robust to varying parameters. For instance, when the gap size was 1 μm, the distance to the truth was 1.3±0.4 μm, the precision was 90±6%, and the sensitivity was 87±15% on average for all the 58 sets of parameters tested.

In our experiments, we applied NeuronStitcher to stitch two images of a mouse V1 neuron from confocal microscopy and another two biocytin-filled human neurons imaged by bright field microscopy (Supplement, Section 4.2, Table S 2). In total, 16 pairs of sections were stitched (the results of 3 adjacent pairs in Figure 1 and more results in Figures S 11-15). Among all automatically matched neurites, 356 (86.6%) were accepted by an expert (Table S 3). NeuronStitcher typically finished the computation within seconds, requiring less than 100Mb memory. The time for a user to visually check and adjust results depended on the complexity of the reconstruction. For our testing datasets, the average time for stitching (including automated computation, visual inspection, parameter fine-tuning, and manually adjustment of the result) an adjacent pair of serial sections was 13’08” (median: 9’41”, minimum: 0’ 13”, and maximum: 36’31”) (Table S 3).

The quality of neuron reconstructions and the selected parameters might influence the performance of NeuronStitcher. To broaden the utility of NeuronStitcher to work with a variety of data acquisition processes, we designed an interactive graphical user interface to (1) allow a visual evaluation on the stitching results and live adjustments of matching parameters; and (2) enable manual corrections of incorrect matching results (Supplement, Section 3). The software was implemented in C/C++ as a plugin of Vaa3D^13,14^, which is a publicly available Open Source platform with a user-friendly interface for 3D+ image analysis and visualization (http://www.vaa3d.org). The screenshot of the GUI and the guidance of the tool can be found in Supplement, Section 3, Figures S 6-9, and Videos S 1-4.

## Acknowledgement

This work was sponsored by the Allen Institute for Brain Science. The authors wish to thank the Allen Institute founders, P. G. Allen and J. Patton, for their vision, encouragement, and support. We thank Zhi Zhou, Staci Sorensen, Lu Li, and Hongkui Zeng for assistance in providing data and discussion of the usability of the software.

